# Organization and dynamics of the SpoVAEa protein, and its surrounding inner membrane lipids upon germination of *Bacillus subtilis* spores

**DOI:** 10.1101/2021.11.20.469378

**Authors:** Juan Wen, Norbert O.E. Vischer, Arend L. de Vos, Erik. M. M. Manders, Peter Setlow, Stanley Brul

## Abstract

The SpoVA proteins make up a channel in the inner membrane (IM) of *B. subtilis* spore. This channel responds to signals from activated germinant receptors (GRs), and allows release of Ca^2+^-DPA from the spore core during germination. In the current work, we studied the location and dynamics of SpoVAEa in dormant spores. Notably, the SpoVAEa-SGFP2 proteins were present in a single spot in spores, similar to the complex formed by all GRs. However, while the GRs’ spot remains in one location, the SpoVAEa-SGFP2 spot in the IM moved randomly with high frequency. The dynamics of the SpoVAEa-SGFP2 and its surrounding IM region as stained by fluorescent dyes were also tracked during spore germination, as the dormant spore IM appeared to have an immobile germination related functional microdomain. This microdomain disappeared around the time of appearance of a germinated spore, the loss of fluorescence of the IM by fluorescent dyes, as well as the appearance of SpoVAEa-SGFP2 peak fluorescent intensity occurred in parallel. These observed events were highly related to the rapid phase darkening, which is considered as the Ca^2+^DPA rapid release. We also tested the response of SpoVAEa and the IM to thermal treatments at 40-80°C. Heat treatment triggered an increase of green autofluorescence, which is speculated to be due to coat protein denaturation, and 80°C treatments induce the appearance of phase-grey-like spores. These spores presumably have a similar intracellular physical state as the phase grey spores detected in the germination but lack the functional proteins for further germination events.

## Introduction

*Bacillus subtilis*, the model Gram-positive bacterium, has multiple complex responses to environmental stress and nutrient depletion. Notably, it can generate different subsets of cells, such as persisters, spores, and biofilms, to promote survival through harsh environmental conditions [1]. Spores in particular are capable of maintaining metabolic dormancy for very long times by protecting their chromosomal DNA through its location in the low water environment of the spore core and its surrounding by multiple protective macromolecular layers [2]. From the outside in, these spore layers are the coat, outer membrane (OM), peptidoglycan (PG) cortex, germ cell wall, inner membrane (IM) and core [2]. Despite their potential long period of dormancy, these spores can be revived or germinated by many environmental cues, primarily small molecules termed germinants that signal the presence of a growth friendly environment, and they then resume cell growth [2].

The SpoVA channel is located in the IM, and has a crucial role in spore germination [3–7]. This channel functions to release the large pool of the 1:1 complex of Ca^2+^ and dipicolinic acid (Ca^2+^-DPA) from the spore core during germination [5,8]. Upon release of this Ca^2+^-DPA, water is taken up and hydrolysis of the PG cortex begins. When the latter is complete the core water content has returned to the levels observed in vegetative cells [2,8]. Multiple signals can trigger the opening of the SpoVA channel, including activated germinant receptors (GRs), hydrolysis of the cortex PG and high hydrostatic pressure [2]. The SpoVA channel has seven subunits, A, B, C, D, Eb, Ea and F [6,9]. SpoVAA, -B, -C, -Eb and -F are transmembrane IM proteins, one of which, SpoVAC, is a pressure-sensitive membrane channel protein [6,7,10,11]. SpoVAD and -Ea are hydrophilic proteins, located outside of the IM, and likely have physical contact with GRs for signal transduction [6,12]. *B. subtilis* spores have an estimated level of ∼7000 molecules of SpoVA proteins, which is 7-10 times higher than the level of GRs [13]. Notably, all GRs in spores are in a complex in the IM that is generally present in only one or two spots/spore termed germinosomes [14–17]. In contrast, by averaging hundreds of consecutive images, previous work indicated that SpoVAEa seems uniformly distributed throughout the IM [17]. In the current work, we created a SpoVAEa-SGFP2 reporter protein. The fusion protein was expressed in *B. subtilis* from the native *spoVAEa* locus and super resolution rescan confocal microscopy (RCM) was used to analyze the SpoVAEa-SGFP2 location in spores.

As mentioned above, GRs are present in foci in the dormant spore IM, and our previous work found that *B. subtilis* spores’ GerKB-SGFP2 foci reached a peak fluorescence intensity around the ‘time to germination’, followed by the dispersion of the spots in a short time window in germinated spores [18]. The ‘time to germination’ was defined as the time needed for the spore to complete half of its rapid decline in phase brightness. The rapid decline of phase brightness is due to the refractility change of spores induced by Ca^2+^DPA release and water uptake and results in the change of a phase bright spore (dormant spore) into a phase dark spore (germinated spore) upon examination under a phase contrast microscope. The molecular basis of GR functioning in spore germination is still not resolved. Here, we speculated that the increased GerKB-SGFP2 fluorescence is due to a relatively ‘dramatic’ environmental change in the physical state of the IM near the fusion protein [18]. Because both GRs and SpoVA proteins are IM proteins, we were curious to assess the dynamic response of SpoVAEa-SGFP2 fluorescence to the environmental changes occurring during germination. Time-lapse imaging was used to track both the refractility change of spores, and the dynamics of SpoVAEa-SGFP2 fluorescence during GR-triggered germination using phase contrast and widefield microscopy. The IM is undergoing dramatic modifications during spore germination, including a 1.3 fold increase in its encompassed volume and the restoration of its lipid mobility [2,19]. Consequently, we also tracked changes in the IM during germination. In addition, because previous work (chapter 4 of this thesis) showed that 40-80°C heat treatments of *B. subtilis* dormant spores resulted in spore heat activation of germination at temperatures ≤ 65°C, caused a combination of heat activation and heat damage at 70-75°C, and led to full heat inactivation at 80°C, we studied thermal effects on SpoVAEa-SGFP2 and membrane dye related fluorescence phenomena.

## Results

### 1. Location and the dynamics of SpoVAEa protein in dormant spores

By using widefield microscopy, we found that the spore surface area did not have a uniform level of SpoVAEa-SGFP2. Next, we employed RCM with a scanning time of 2 sec per frame to obtain more detailed structural information [20]. Surprisingly, clusters of GFP fluorescence were observed, which clearly stood out from the background in dormant PS832 SpoVAEa-SGFP2 spores (**Fig. 1A, left panel**). By enlarging one of the spores, two spots were observed in the spore labelled ‘a’ (**Fig. 1A, right panel**). The full width at half maximum (FWHM) of the pronounced spot of spore ‘a’ was 220 nm (**Fig. 1C**). Due to the fact that the visualization of germination proteins can be interfered with by heavy autofluorescence from the spore coat, we also imaged here SpoVAEa-SGFP2 in a coat defective background [21]. Similar fluorescent spot structures were found in the coat defective PS4150 SpoVAEa-SGFP2 spores (**Fig. 1B, C**). These spots were, however, not more pronounced indicating that our observations in wild-type spores were not majorly obscured by coat layers.

**Figure 1.**
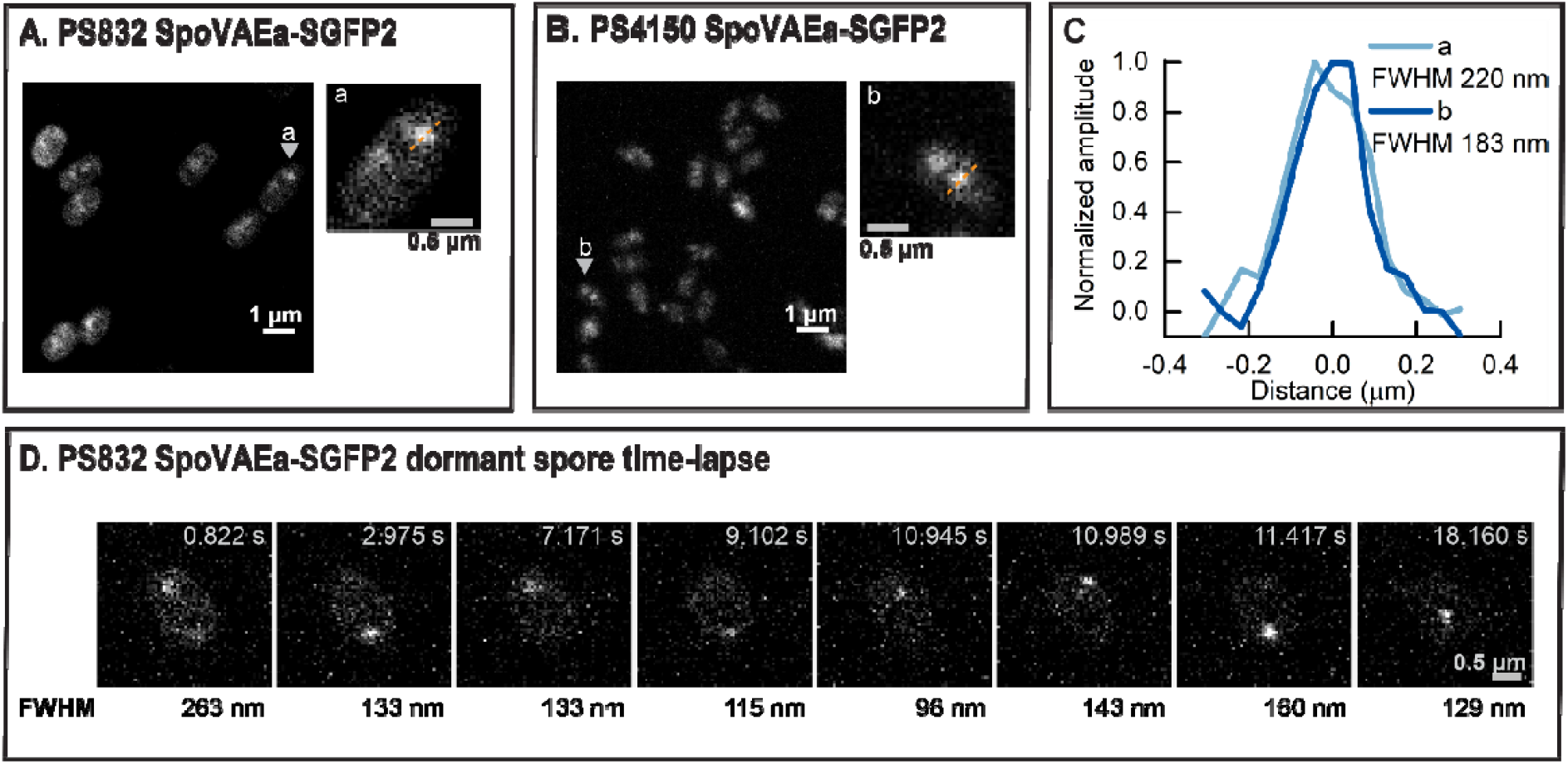
Organization of SpoVAEa-SGFP2 in dormant spores in a region of increased fluorescence (a fluorescent ‘spot’). (A) The RCM fluorescence image of PS832 SpoVAEa-SGFP2 spores. The spore in the inset ‘a’ is the enlarged view of the spore indicated by the grey arrow in the left panel. (B) The RCM fluorescence image of PS4150 SpoVAEa-SGFP2 spores. The spore in the inset ‘b’ is the enlarged view of the spore indicated by the grey arrow in the left panel. (C) The plot profiles of the pronounced SpoVAEa-SGFP2 spot of spores ‘a’ and ‘b’. (D) The RCM high frequency time lapse images (22.2 ms/frame) of a PS832 SpoVAEa-SGFP2 dormant spore. The presented images represent frames in which the strong SpoVAEa-SGFP2 spot appears during the high frequency time lapse track. The relative FWHM of the SpoVAEa-SGFP2 spot in each frame is indicated at the bottom of each image.

Previously, we showed that GerKB-SGFP2 proteins, one class of GRs, presented themselves as spots [18]. These spots ‘blinked’ within the confinement of a small area of the IM, at high frequency [18]. Here, the crucial question is whether SpoVAEa-SGFP2 is also at least somewhat mobile in the IM. To address this query, RCM microscopy was applied for high frequency imaging with a scanning time of 22.2 milliseconds per frame. Even more surprisingly, a single spot, traveling around the IM, was observed in the PS832 SpoVAEa-SGFP2 dormant spores, and the spore was tracked for 1000 frames. We presume that we are the first to see this movement as we imaged at very high time resolution. The fluorescent spot of the spore ‘randomly’ appeared in different time frames and different locations of the spore. **Fig. 1D** shows eight frames with a pronounced fluorescent spot, and the location differs in each frame. The measured FWHM of the spot in different frames varied from 96 nm to 263 nm. Hence, the observed high frequency movement with longer exposure time, could explain why not all the spores in **Fig. 1A** and **Fig. 1B**, showed a spot structure, as well as the uniform distribution of SpoVAE in previous work [17]. It is, however, not clear how SpoVAEa is able to move at such a high frequency within certain boundaries in an otherwise ‘rigid’ inner membrane [19,22]. Moreover, we cannot exclude that there are ‘free’ SpoVAEa proteins distributed over the IM outside of the spot.

### 2. Dynamics of SpoVAEa in the IM of *B. subtilis* spores during GR-triggered spore germination

Previously, we found that during spore germination GerKB-SGFP2 spots gradually increased in fluorescence intensity [18]. This phenomenon is potentially related to the change of the spore’s physical state in general and the IM in particular upon germination, because no new protein synthesis was observed [18]. It could be linked, for instance, to the increase in core water content and core pH due to the release to Ca^2+^-DPA. Here we used widefield microscopy to track the overall mean intensity of SpoVAEa-SGFP2 in spores during germination via GerB and GerK GRs by supplying the nutrient germinant cocktail AGFK (L-asparagine, glucose, fructose, and potassium chloride).

The phase brightness and SpoVAEa-SGFP2 fluorescence history of a single spore is shown in the time-lapse image montage in **Fig. 2A and 2C**. By analysing the brightness and fluorescence profiles of this spore, we observed that the peak fluorescence intensity of SpoVAEa-SGFP2 was reached before the appearance of the phase dark spore, followed by a slow decline of fluorescence intensity in the phase dark spore (**Fig. 2A-D**). The initiation of SpoVAEa-SGFP2 fluorescence increase was at the same time as the start of the spore’s brightness rapid decline, which is considered as the start of Ca^2+^-DPA rapid release (**Fig. 2B, D, E**). In order to confirm the observed dynamics in the population, we synchronized SpoVAEa-SGFPs fluorescence profiles by defining the t=0 as the ‘time to germination’, which is the time needed for the spore to complete half of its rapid decline in phase brightness (**Fig. 2E**). In the averaged trace of 418 germinating spores, the SpoVAEa-SGFP2 intensity increased sharply and the peak was around the ‘time to germination’ (**Fig. 2F**), while no SpoVAEa-SGFP2 synthesis was detected by western blot analysis (**Fig. S1**). Thus, the increase of SpoVAEa-SGFP2 fluorescence intensity during germination is correlated with the rapid release of Ca^2+^-DPA. Western blot analysis in the current work, as well as previous works, did not detect a significant decrease of SpoVAEa level in germinated spores compared to dormant spores (**Fig. S1**), however, a decrease of SpoVAEa-SGFP2 fluorescence was observed in germinated spores (**Fig. 2D, 2F**) [6]. Subsequently, we found that photobleaching had a role in the observed fluorescence decrease (**Fig. S2**), and photochemical alteration of both spore coat and SGFP2 potentially contributed to the photobleaching. A spore coat defective strain with eliminated autofluorescence will likely be necessary to study the dynamics of SpoVAEa-SGFP2 in germinated spores [21,23].

**Figure 2.**
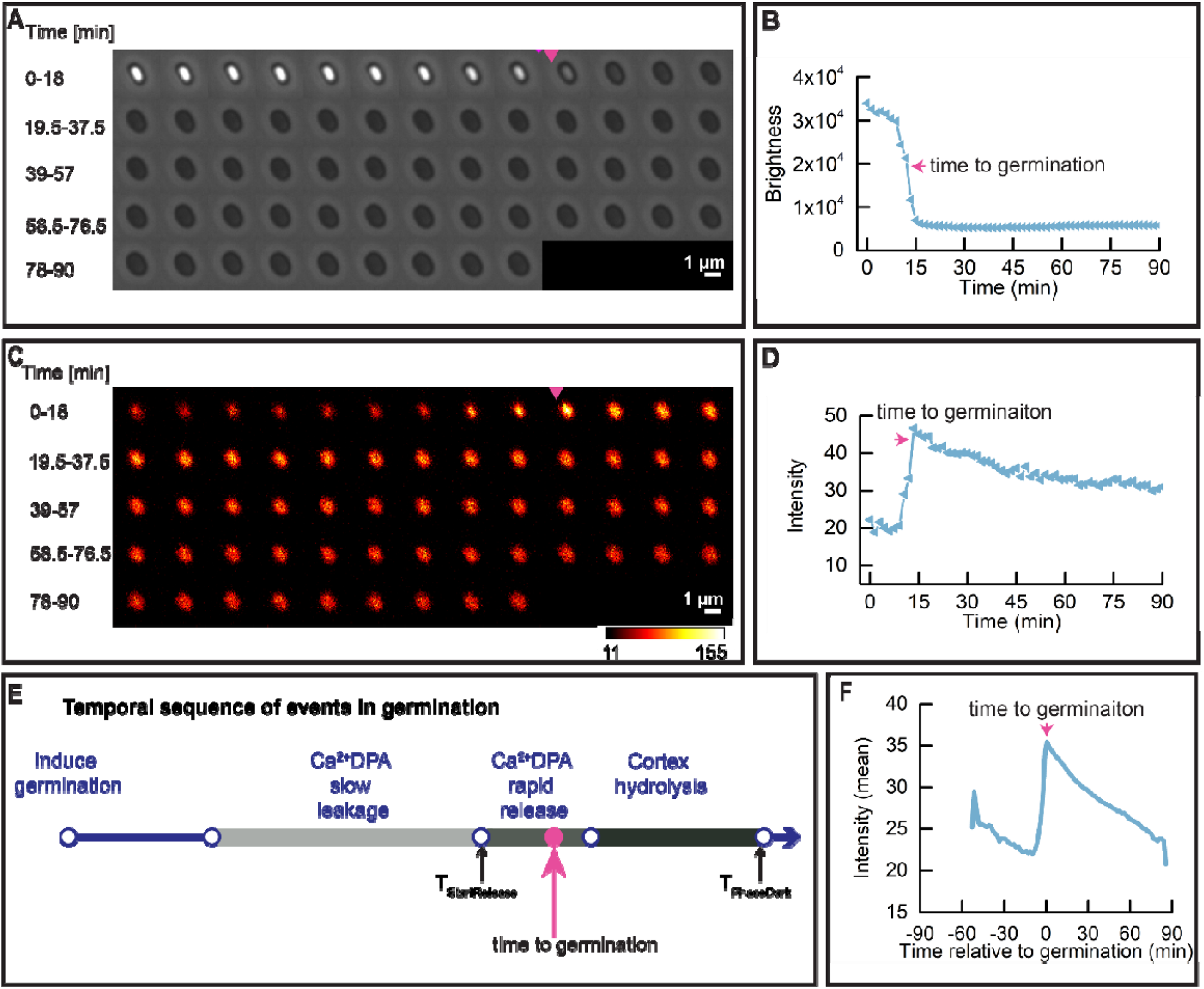
Dynamic behaviour of SpoVAEa-SGFP2 during spore germination. Spore germination was triggered by (10 mM each) AGFK without heat activation treatment. (A) Phase contrast time lapse images of a single PS832 SpoVAEa-SGFP2 spore. (B) The brightness profile corresponding to the images shown in panel A. (C) The fluorescence time lapse images of the same spore shown in panel A. (D) The SpoVAEa-SGFP2 intensity profile corresponding to images in panel C. (E). Parameters detected by SporeTrackerX, and their corresponding positions in spore germination. As shown in the schematic, upon germination commitment, Ca^2+^DPA starts slow leakage from the spore core, followed by the rapid release of the remaining Ca^2+^DPA and then spore cortex hydrolysis [24]. The latter two events result in the rapid decline in the brightness of spore in phase contrast microscope. Here, SporeTrackerB_06K detected the T_StartRelease_ (time of initiation of the rapid decline in spore brightness) and T_PhaseDark_ (time of completion of spore brightness rapid decline) in the brightness profile, and further calculated the ‘time to germination’ for presenting data for the population. Here, the ‘time to germination’ is defined as the time needed for the spore to complete half of its rapid decline in phase brightness. The magenta arrow indicates the ‘time to germination’. (F) Average of 418 synchronized single SpoVAEa-SGFP2 spore intensity traces. Synchronization defines t=0 min as the ‘time to germination’. 613 spores’ were tracked by microscopy for 90 min and 68.2% of them completed germination.

### 3. Changes in IM structure upon triggering spore germination via GRs

It is known that lipids in the *B. subtilis* dormant spores are immobile [2,19]. Still, this IM is capable of its increasing surface area ∼1.3 fold upon the spore swelling due to spore core water uptake and cortex hydrolysis [2]. As indicated above, SpoVAEa-SGFP2 fluorescence reached peak intensity around ‘time to germination’. Consequently, we decided to probe for a putative correlation between the SpoVAEa-SGFP2 peak fluorescence intensity and the change in mobility of the IM lipids. To that end, the IM of PS832 spores was stained with either the carbocyanine dye DiIC_12_ or the styryl dye FM5-95, which were added to a sporulating culture and hence incorporated into the IM upon the formation of the forespore, as has been shown in previous studies [16,19]. Any lipid probe present in the OM could be removed by extensive washing during spore purification [19].

We again used widefield microscopy to track the change in IM staining during germination. The fluorescence intensity of DiIC_12_ and FM5-95 stained spores dramatically dropped upon the start of the spore’s brightness rapid decline, and the drop was completed around the ‘time to germination’ (**Fig. 3, 4**). In a word, just as with the dynamics of SpoVAEa-SGFP2, the change of the IM during germination is also highly correlated with the Ca^2+^-DPA rapid release, which leads to the increase of spore core pH, and water content [25,26]. The FM5-95 stained spore also had a fluorescent spot, whereas DiIC_12_ spores had relative uniform staining (**Fig. 3C, 4C**). Regarding the fact that the FM5-95 dye was almost invisible in the germinated spore (**Fig. 4C**), we note that the hydrophobicity and affinity of FM5-95 for the IM lipids seems lower than that of DiIC12. This implies also that FM5-95 prefers to bind into lipid domains with higher fluidity. Hence, the FM 5-95 spot area could well be a fluid membrane domain, instead of the compressed IM. The disappearance of the FM5-95 spot in the phase dark spore, suggests that this region might be a germination related functional membrane microdomain (**Fig. 4C**).

**Figure 3.**
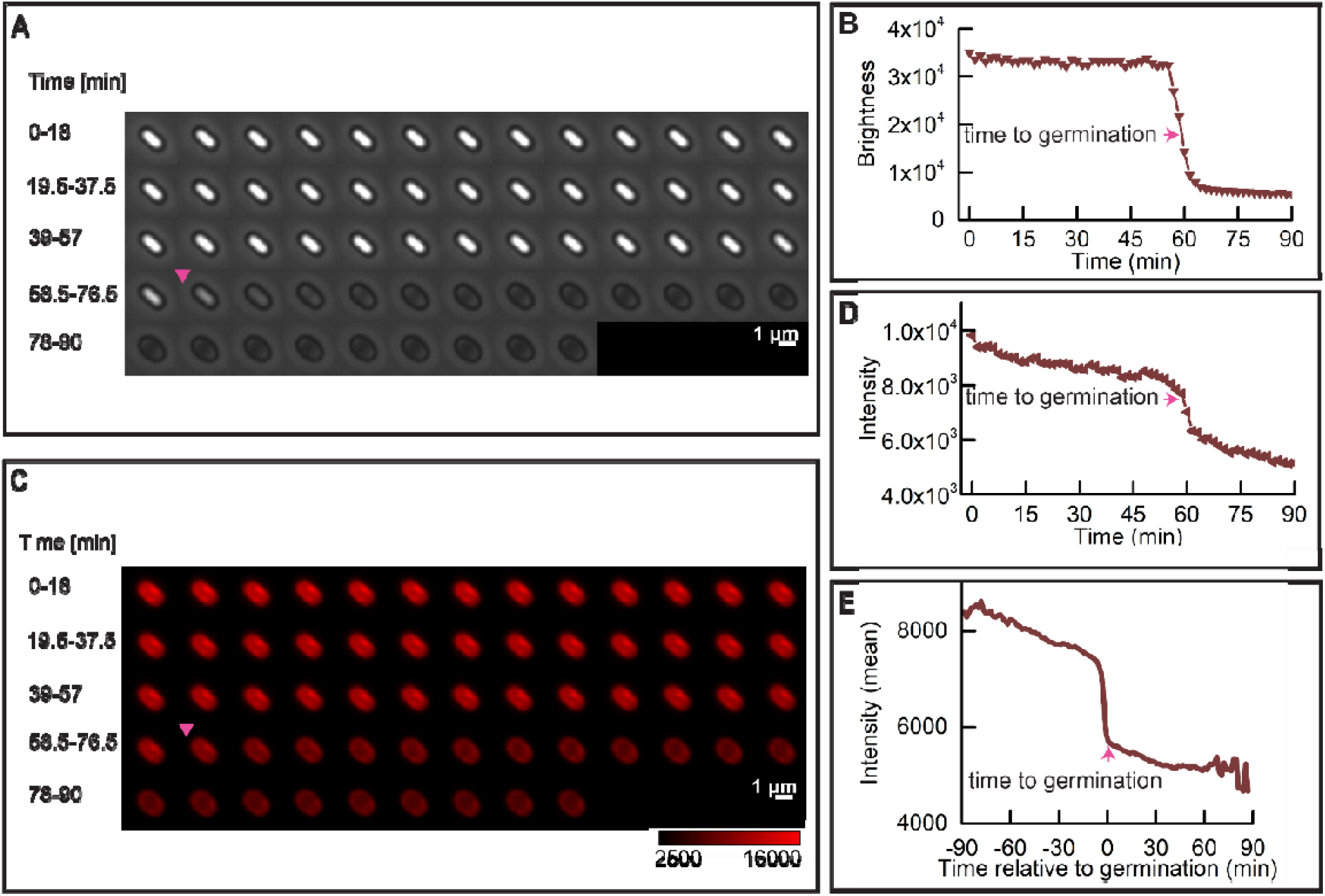
Dynamic behavior of the DiIC_12_ stained wild-type spore IM during germination. Spore germination was triggered by (10 mM each) AGFK as described in Methods. (A) Phase brightness of a PS832 (DiIC_12_) spore. (B) The spore brightness profile corresponding to the images shown in panel A. (C) The fluorescence time lapse images of the same spore shown in panel A. (D) The DiIC_12_ stained IM intensity profile corresponding to images in panel C. The magenta arrow indicates the ‘time to germination’. (E) Average of 170 synchronized single DiIC_12_ spore fluorescence intensity traces. Synchronization defines t=0 min as the ‘time to germination’. 289 spores’ were tracked by microscopy for 90min, and 58.8% of them completed germination.

**Figure 4.**
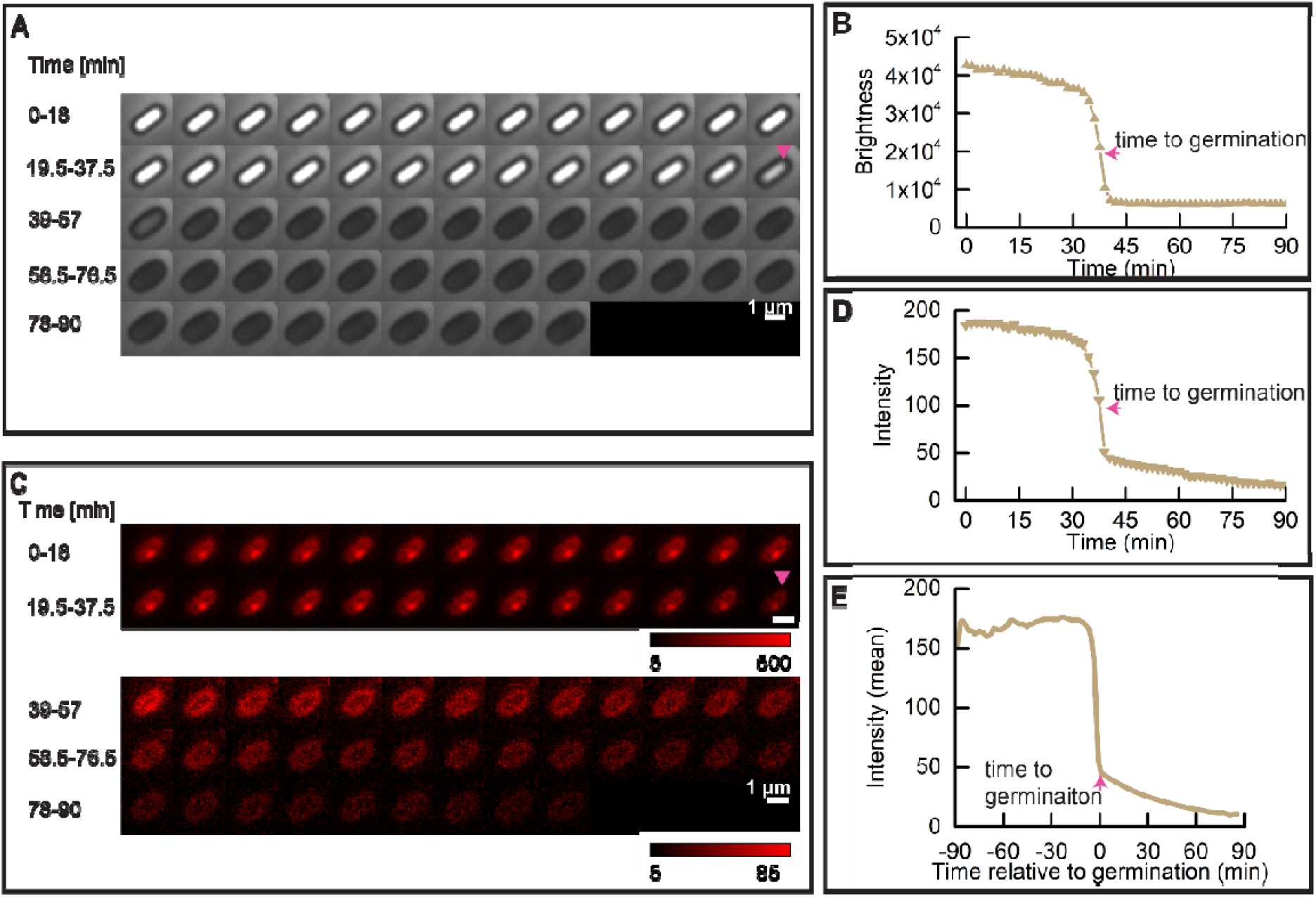
Dynamic behavior of the FM5-95 stained IM during wild-type spore germination. Spore germination was triggered by (10 mM each) AGFK as described in Methods. (A) Phase brightness of a PS832 (FM5-95) spore. (B) The spore brightness profile corresponding to the images shown in panel A. (C) The fluorescence time lapse images of the same spore shown in panel A. (D) The FM5-95 stained IM intensity profile corresponding to images in the panel C. The magenta arrow indicates the ‘time to germination’. (E) Average of 277 synchronized single FM5-95 spore fluorescence intensity traces. Synchronization defines t=0 min as the ‘time to germination’. 369 spores’ were tracked by microscope for 90min, and 75.1% of them completed germination.

### 4. Response of SpoVAEa and probe stained IM to thermal treatments

Our previous work showed that heating *B. subtilis* spores at 40-65°C promotes spore germination in a positive time/temperature dependent manner, higher heat temperatures resulted in both heat activation and heat damage, and eventually led to heat inactivation at 80°C (chapter 4 of this thesis). The germination proteins and their surrounding IM are potential targets of the heat treatments. Here we studied whether heat treatments change the states of the spore’s, SpoVAEa and the IM after 5 hours of heat treatment at 40-80°C.

Significant drops of spore’s absorbance at the population (**Fig. 5A, 7A, 7E**) and refractive index at single spore level (**Fig. 5C, 5E, 7C, 7G**) were detected in spores treated at 80°C, and in some cases also in 75 °C treated spores. Spores treated at 40-70°C morphologically looked similar upon the microscopical examination. Hence, we only present images of 65, 75, and 80°C heated spores as representative images (**Fig. 6, 8**). As shown in the phase contrast images (**Fig. 6, 8**), a subpopulation of phase-grey-like spores appeared in 80°C treated groups. These phase-grey-like spores are potentially spores that have lost Ca^2+^-DPA, as reported previously [27,28].

**Figure 5.**
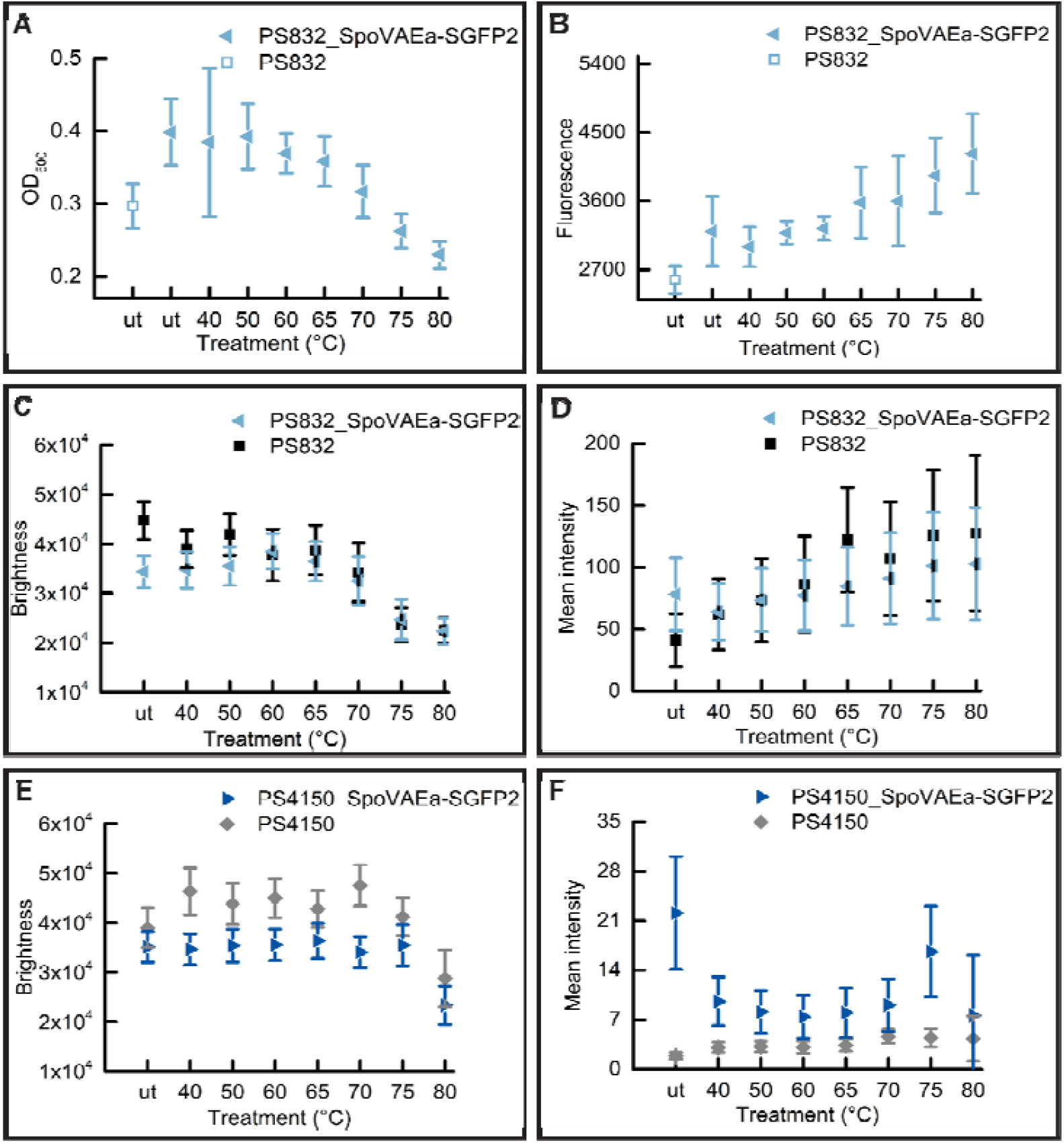
Changes in SpoVAEa-SGFP2 dormant spore refractility and fluorescence intensity at population and single spore levels after 5 hours of heat treatment at various temperatures. (A) The change of OD at 600nm of a PS832 SpoVAEa-SGFP2 dormant spore population measured by a plate reader. Untreated PS832 spores were used as control. (B) The corresponding fluorescence intensity of spores in panel A. (C) The change of brightness of PS832 SpoVAEa-SGFP2 dormant spores at single spore level measured by a phase contrast microscopy. Here, PS832 spores with different heat treatments were employed as control. ≥ 444 spores were examined in each group. (D) The corresponding intensity of spores in panel C measured by widefield microscopy. (E) The brightness of PS4150 SpoVAEa-SGFP2 dormant spores at single spore level. Here, PS4150 spores with different treatments were employed as control. ≥ 937 spores were examined in each group. (F) The corresponding fluorescence intensity of spores in panel E. The number of spores detected by microscopy is presented in Table. S1.

**Figure 6.**
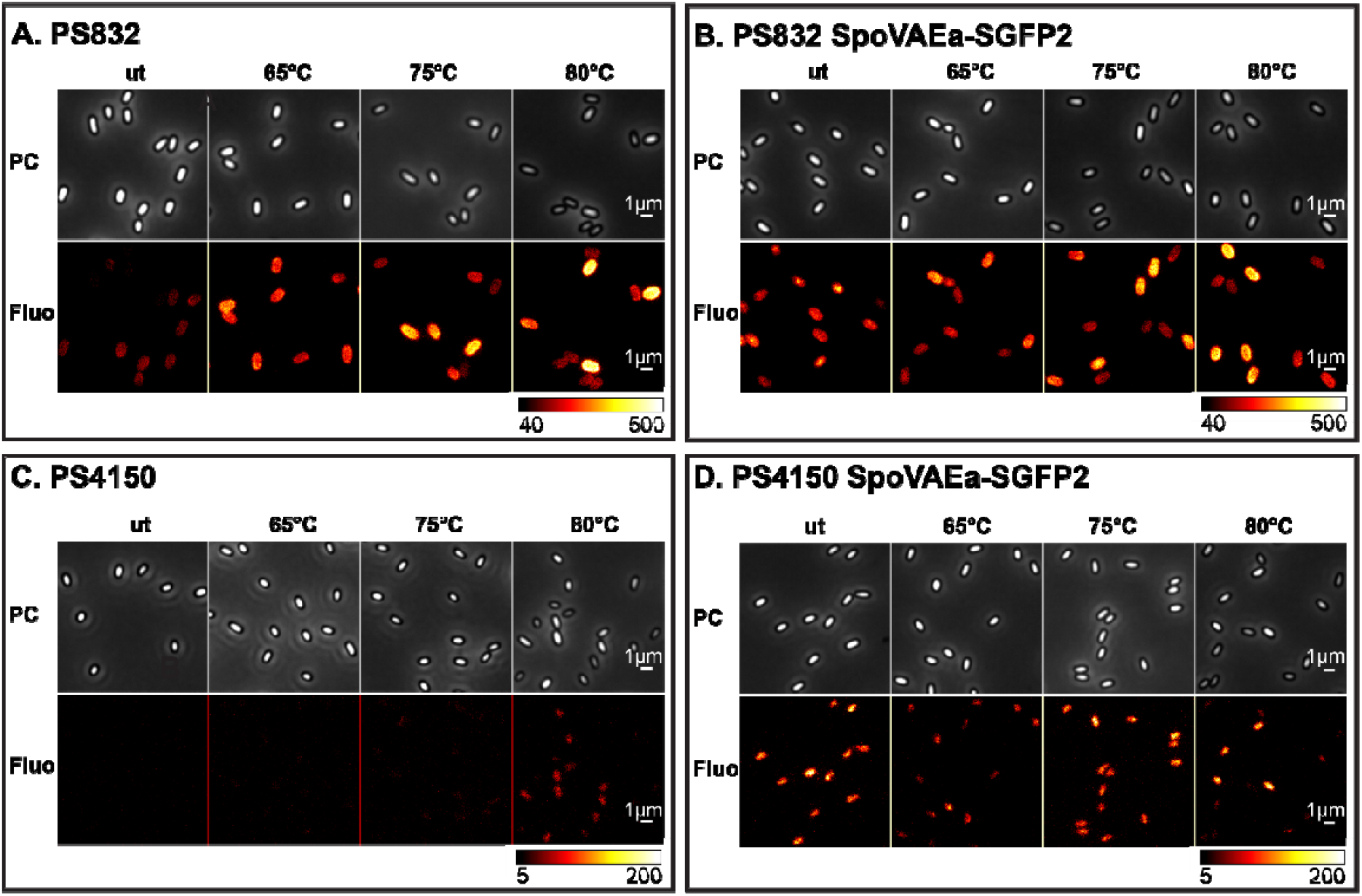
The effect of 5 hours of heat-treatment on PS832 SpoVAEa-SGFP2. The phase contrast (PC) and widefield fluorescence (Fluo) images of spores of wild type strain PS832 (A), PS832 SpoVAEa-SGFP2 spores (B), (C) PS4150, and (D) PS4150 SpoVAEa-SGFP2 after different heat treatments. (A) and (B) are the images corresponding to Fig. 5C-5D. (B) and (C) are the images corresponding to Fig. 5E-5F. The fluorescent image intensity display range for each strain is the same.

**Figure 7.**
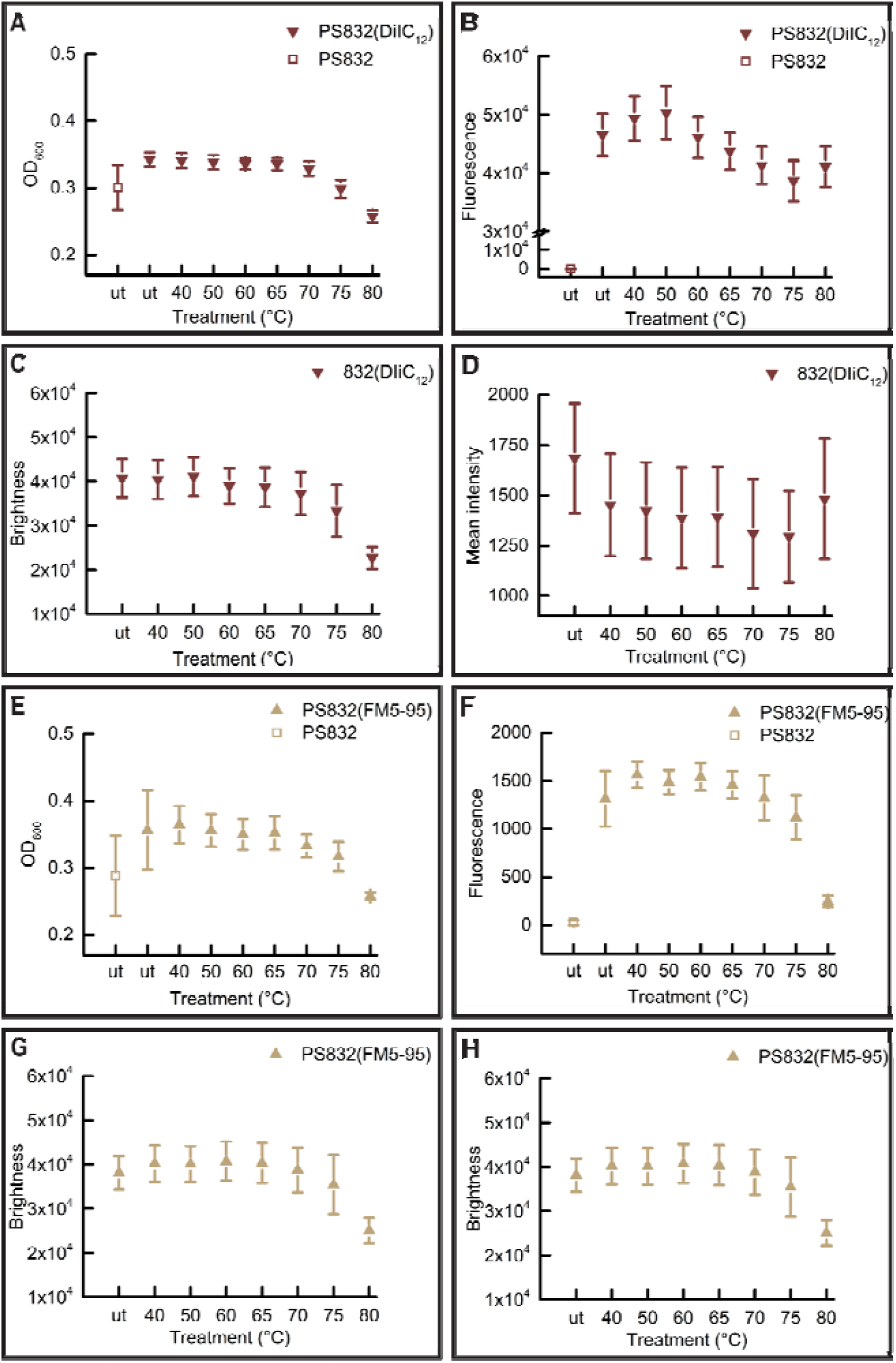
The change of lipid probe stained dormant spore refractility and fluorescence intensity at population and single spore levels after 5 hours of treatment at various temperatures. (A) The change of OD_600_ of PS832 (DiIC_12_) dormant spore populations measured by a plate reader. Untreated PS832 spores were used as control. (B) The corresponding fluorescence intensity of spores in panel A. (C) The change of individual PS832 (DiIC_12_) spores’ brightness measured by a phase contrast microscopy. ≥ 612 spores were examined in each group. (D) The corresponding fluorescence intensity of spores in panel C. (E) The brightness of individual PS832 (FM5-95) dormant spores. ≥ 932 spores were examined in each group. (F) The corresponding fluorescence intensity of spores in panel E. The numbers of individual spores examined by microscopy are given in Table. S2.

**Figure 8.**
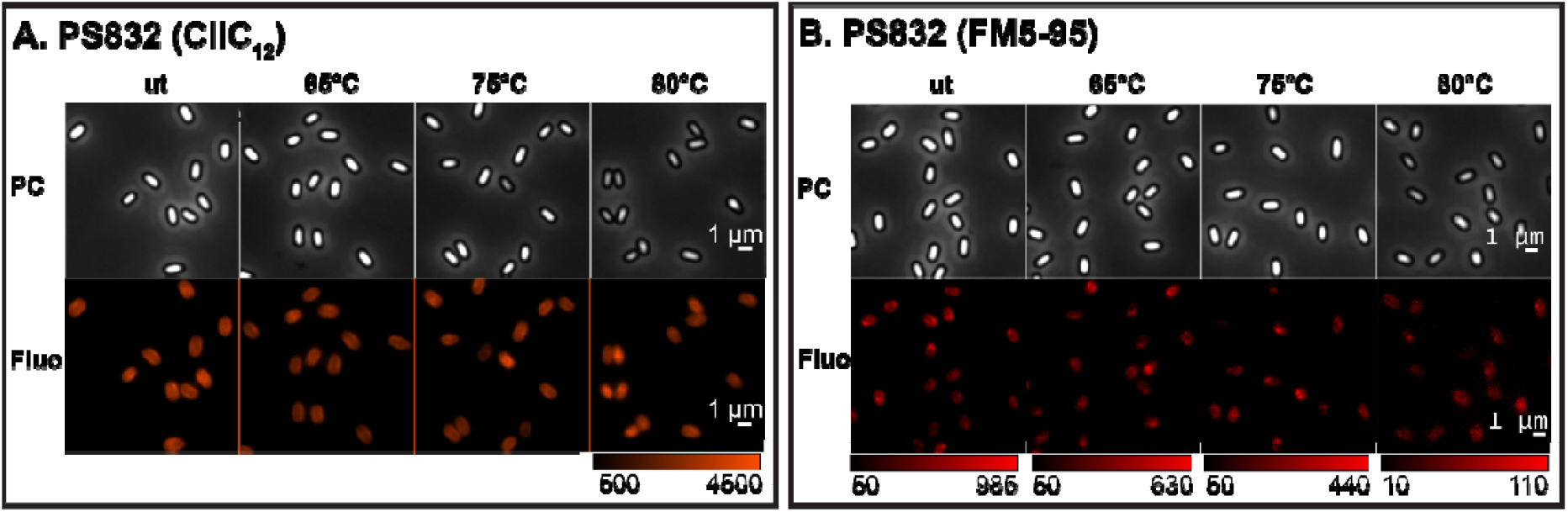
The effect of 5 hours of heat-treatment on the fluorescence intensity of probe stained spores. The phase contrast (PC) and widefield fluorescence (Fluo) images of wild type PS832 spores stained by DiIC_12_ (A), and FM5-95 (B) after different heat treatments: (A) images corresponding to Fig. 7C-7D; and (B) images corresponding to Fig. 7G-7H. The fluorescence images displayed for DiIC_12_ spores are all in the same range, whereas, the FM5-95 fluorescence images displayed are in different ranges due to the big decline in brightness at higher temperatures.

Heat treatments at 40-80°C led to a decrease of fluorescence intensity of coat defective PS4150 SpoVAEa-SGFP2 spores (**Fig. 5F, 6D**). However, we cannot exclude the possibility that the decreased SpoVAEa-SGFP2 spore fluorescence was caused by heat denaturation of SGFP2 [29]. This denaturation seems reversible at temperatures ≤65°C, due to the fact that 65°C treated PS832 SpoVAEa-SGFP2 spores still exhibited similar fluorescence profiles as untreated spores during spore germination (**Fig. S2, 2**). However, it is not clear how spores maintain functionally active channel proteins during the heat activation process. Clearly, a pH and thermal stable fluorescent reporter will be important for future spore related research [30–32]. Notably, we tested the response of SpoVAEa-SGFP2 to heat in coat defective spores, because the change of SpoVAEa-SGFP2 fluorescence was masked by the enhanced green autofluorescence of the spore coat (**Fig. 5, 6**). The increase of autofluorescence had a positive correlation with the heat treatment temperature. We speculate the enhanced autofluorescence is likely related to coat protein denaturation, since studies showed that a lethal thermal treatment induced significant protein denaturation in *B. subtilis, B. cereus, and B. megaterium* spores [27,28]. Besides, the observed increased autofluorescence in PS4510 spores treated at 80°C is considered due to the response of either the residual outer coat or the inner coat layers to heat (**Fig. 6C**).

Compared to the untreated (ut) DiIC_12_ labeled spores, spores heated at 40-80°C exhibited slightly decreased fluorescence intensity (**Fig.7B, 7D, 8A**). However, no correlation was detected between the drop of the phase contrast image brightness and changes in DiIC_12_ fluorescence (**Table.S2**). In contrast, FM5-95 stained spores showed a continuous fluorescence decrease upon treatment at 70, 75, and 80°C (**Fig.7F, 7H, 8B**). The FM5-95 fluorescence had a positive correlation with the spore brightness of 70 and 75°C heated spores (**Table.S2**), Pearson correlation coefficient 0.54 and 0.76, respectively, and it was almost invisible in spores heated at 80°C. As mentioned above, we speculate that spores with decreased brightness are potentially spores that have lost Ca^2+^-DPA. Hence, the decrease of FM5-95 intensity presumably correlated with the release of Ca^2+^-DPA, and the subsequent physical state changes in spores.

## Discussion

Our work shows that at least one SpoVA subunit SpoVAEa, was clustering as a spot undergoing high frequent movement in dormant *B. subtilis* spore. In addition, a germination related microdomain with higher fluidity was observed in the spore IM. The rapid phase darkening of IM labeled spores, caused by the rapid release of Ca^2+^DPA and cortex hydrolysis, is accompanied by the loss of fluorescence in the IM the disappearance of the IM fluid microdomain, and an increase in SpoVAEa-SGFP2 intensity. Heat treatment at 65-80°C resulted in a temperature dependent increase in green autofluorescence, potentially caused by the degradation of coat proteins. Dormant spores heat treated at 80°C have a sub-population of phase-grey-liked spores.

As mentioned above, a FM5-95 stained microdomain was observed in dormant spores, and this domain disappeared upon the appearance of the phase dark spore (germinated spore). In addition, the microdomain likely has higher fluidity compared to the rest of the IM, and remains in a confined location in the IM before dispersing upon germination. This immobile behavior is in line with the activity of the ‘germinosome’ spot, in which GRs and their scaffold protein GerD are clustered together [17]. In addition, our previous work showed that the ‘germinosome’ agglomerates and diffuses in the same location of the IM, and gradually disperses after spore phase transition [15,18]. Hence, there is a potential link between the germinosome and the fluid microdomain. To better understand this germination related membrane domain, efforts can take aim at answering the following questions. 1) Is the ‘germinosome’ spot truly located in the microdomain? 2) Does GerD have a function in recruiting the specific disordered lipids into this microdomain, as it does play such a role in maintaining the ‘germinosome’ spot? 3) When and how does the fluid microdomain form during sporulation?

Except for the microdomain in the IM, the spore core membrane itself still remains mysterious. Our current work, as well as previous publications showed that there is an expansion process of the IM surrounded area and a reduced staining by lipid dyes in the IM during germination [19]. Notably, it has long been known that this dramatic increase of the IM surrounded area takes place without new lipid synthesis, and a recent electron microscopy study revealed an intracellular membrane structure in dormant spores below the IM [19,33]. This membrane structure, which contains at least one SpoVA protein, disappears upon spore core hydration, most likely due to integration with the IM [33]. The integration of the IM lipids during the germination might have a role in response to the decline of the dye staining in the IM. A technique with better temporal and spatial resolution might show the dramatics of the integration. Current work showed that SpoVAEa clusters in the IM as a spot, and is capable of moving in the ‘gel state’ IM randomly with high frequency [19]. This random movement of the SpoVAEa spot might be beneficial for the physical interaction with the immobile GR spots. However, it is not clear how the SpoVA channel proteins interact with GRs, nor whether this interaction requires the integration of all SpoVA proteins either in the spot or dissociated in the IM, or even from the intracellular membrane structure. In addition, SpoVA channel proteins, synthesized in the developing forespores, are also crucial for import of Ca^2+^-DPA during sporulation [6,34]. Revealing the mechanism of SpoVA spot formation in the forespore will contribute to the knowledge of both Ca^2+^-DPA uptake and release processes.

We also made efforts to check the response of SpoVAEa-SGFP2 and the dye stained IM to heat treatments, which induced either heat activation or heat inactivation to spores. The thermal inactivation resulted in the appearance of phase-grey-like spores. However, no clear changes were observed in spores treated at heat activation temperatures (40-65°C). Interestingly, the green autofluorescence, related to the proteinaceous coat, increased with the elevated heat temperatures. We speculate this phenomenon likely relates to coat protein’ denaturation. Studying spore proteins’ thermal stability profile on the proteome scale might provide a way to understand heat activation and inactivation at the molecular level.

## Materials and Methods

### Bacterial strains, and culture conditions

*B. subtilis* PS832 is a prototrophic 168 laboratory strain. Strain PS4150 is derived from PS832 carries the *cotE* and *gerE* deletion mutations and is a coat defective mutant [21]. Strains PS832 SpoVAEa-SGFP2 and PS4150 SpoVAEa-SGFP2 were derived from PS832 and PS4150, respectively. These two strains contain a SpoVAEa-C-terminal SGFP2 fusion, integrated at the *spoVAEa* locus, and expressed under the control of the *spoVA* operon′s promoter [35]. Spores of all strains were prepared in 2×SG sporulation medium at 37 °C as described previously [36]. If required, the fluorescent dyes DiIC_12_ (1 μg/ml, ThermoFisher) or FM5-95 (2 μg/mL, ThermoFisher) were added to the sporulation culture, when it reached the peak optical density at 600 nm (OD_600_). Spores were harvested and purified, including centrifugation through HistodenZ as described previously [16]. Spores (OD_600_ ∼ 60, in MIlliQ water) used in the current work had ≥98% dormant spores, and were essentially free of germinated spores, cells or cell debris as verified by phase contrast microscopy.

### Measuring the fluorescence of SpoVAEa-SGFP2 and the dye stained IM in heated spore populations

Spores (OD600 = ∼ 1, in MIlliQ water) were heated for 5 h at 40, 50, 60, 65, 70, 75 or 80°C, followed by cooling in a water-ice bath (⍰15 min). Spores (OD_600_ = ∼1, 150 μl/well) were added to a 96-well flat-bottomed microtiter plate (black wall, Greiner Bio-One), and the optical density and fluorescence intensity were measured by a BioTek plate reader. Data were collected from at least two independent tests, each of them had three biological repeats.

### Imaging and image analysis

In current work, two microscopes were employed for imaging. The widefield microscope had a Nikon Ti Microscope, a NA1.45 plan Apo *λ* 100× Oil Ph3 DM objective, and the rescan confocal microscope was equipped with a Nikon Ti Microscope and a NA1.49 SR Apo TIRF 100× objective. Spores were stabilized on a 1.5% agarose pad sealed in an air containing chamber as described previously [37]. If necessary, spores were heated for 5 h at 40, 50, 60, 65, 70, 75 and 80 °C, followed by cooling in a water-ice bath (⍰15 min). In case of tracking spores’ germination, HEPES buffer (25 mM) supplemented with AGFK (L-asparagine, glucose, fructose, and potassium chloride, 10 mM each) were provided in the agarose pad. The time lapse images were captured once every 1.5 min for 90 min.

Time lapse images were analyzed by the ImageJ macro SporeTrackerB_06k [38]. In brief, spores were detected and marked in the first-time frame of the phase contrast images. For each spore, a phase contrast and a fluorescence montage stack were created to present the phase contrast brightness history, and the fluorescent dynamics of this spore (**Fig. 2A, 2C**). The phase contrast brightness profile and fluorescent profile of the spore were detected and stored by SporeTrackerB_06k for display and further analysis (**Fig. 2B, 2D**). Subsequently, the time of start (T_RapidRelease_) and (T_PhaseDark_) end of rapid decline in the brightness profile were detected, and further the ‘time to germination’ was calculated for presenting the averaged fluorescence profile in the population (**Fig. 2E**) [24]. Here, the ‘time to germination’= 1/2 × (T_RapidRelease_+T_PhaseDark_). In order to present the fluorescent profile of the population, each spore’s fluorescence profile was synchronized by defining t=0 min as the ‘time to germination’. Eventually, averaged fluorescence traces vs time relative to germination were created to show the fluorescence profile in the population (**Fig. 2F**). Two channel images (phase contrast and fluorescent image) with single time frames were analyzed by ImageJ macro SporeAnalyzer. Spores were detected and marked in the phase contrast channel, followed by the detection of spore brightness and fluorescence intensity of each spore. Both macro runs in the background of ImageJ plugin ObjectJ.

### Western blot

*B. subtilis* PS832 SpoVAEa-SGFP2 spores were heat activated for 30 min at 70 °C, followed by cooling in a water-ice bath (⍰15 min). Subsequently, spore germination (OD600 = ∼ 30, 100 μl) was triggered by L-asparagine, glucose, fructose, and potassium chloride, 10 mM each (AGFK) in 5 ml MOPS medium. Spores were collected after incubation at 37 °C for 0, 15, 30, and 60 min with continuous rotation at 200 rpm. Spore lysates were obtained by the procedure of Troiano *et al*. [15]. Proteins from equal aliquots of the same amounts of spores were run on a Tricine-SDS-PAGE gel, and probed with rabbit polyclonal anti-GFP antibody (Abcam) and HRP-conjugated goat anti-rabbit IgG H&L (Abcam) on a PDVF membrane [39].

## Supporting information

Supplementary data SpoVAEa B. subtilis

## Acknowledgements

We thank the Van Leeuwenhoek Centre for Advanced Microscopy (LCAM) at the University of Amsterdam for the use of the widefield microscope, and the Confocal.nl for the use of the RCM microscope. We thank Ronald Breedijk from LCAM for the assistance for imaging. We thank Jeroen Kole from Confocal.nl for the RCM imaging. Juan Wen acknowledges the China Scholarship Council for a PhD fellowship.

## Author contributions

Juan Wen (data analysis and interpretation, manuscript writing), Norbert O. E. Vischer (image analysis), Arend D. Vos (data analysis and interpretation), Erik. M. M. Manders (RCM microscopy), Peter Setlow (data interpretation and manuscript editing) and Stanley Brul (data analysis, manuscript interpretation, editing and project supervision).

## Notes

https://hdl.handle.net/11245.1/068d0aaf-4e64-42d3-a808-4ae199823678

### Competing Interest Statement

The authors have declared no competing interest.

### Summary of Updates

Author name of Arend L. de Vos wasn’t spelled correctly.

