## Supplementary data SpoVAEa B. subtilis for "Organization and dynamics of the SpoVAEa protein, and its surrounding inner membrane lipids upon germination of *Bacillus subtilis* spores"

**Supplementary information**

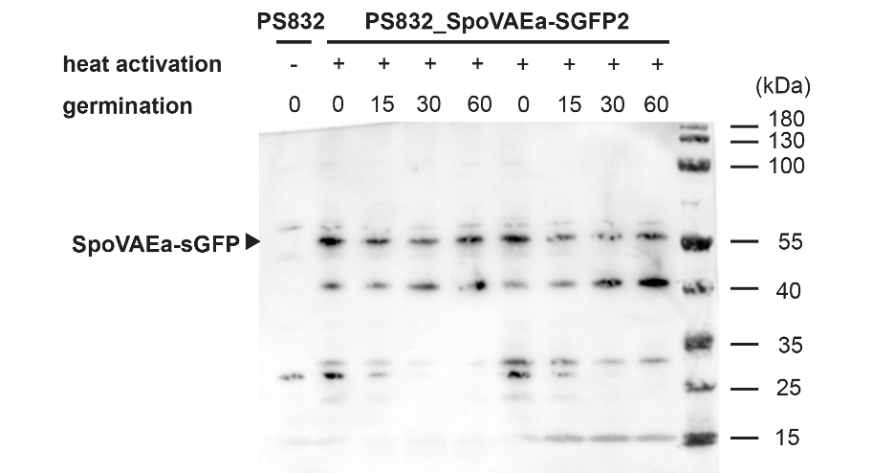

**Figure S1.** Western blot of SpoVAEa-SGFP2 in dormant and germinated PS832 SpoVAEa-SGFP2 spores. Spore germination was induced by (10 mM each) AGFK after a 30 min heat treatment at 70 °C for promoting and synchronizing spore germination. Proteins were extracted from spores germinated for 0, 15, 30, and 60 min, run on SDS-PAGE, proteins transferred to a PDVF membrane and SpoVAEa-SGFP2 was detected with polyclonal rabbit anti-GFP antibodies (Abcam) followed by an HRP-conjugated secondary goat anti-rabbit lgG H&L antibody (Abcam).

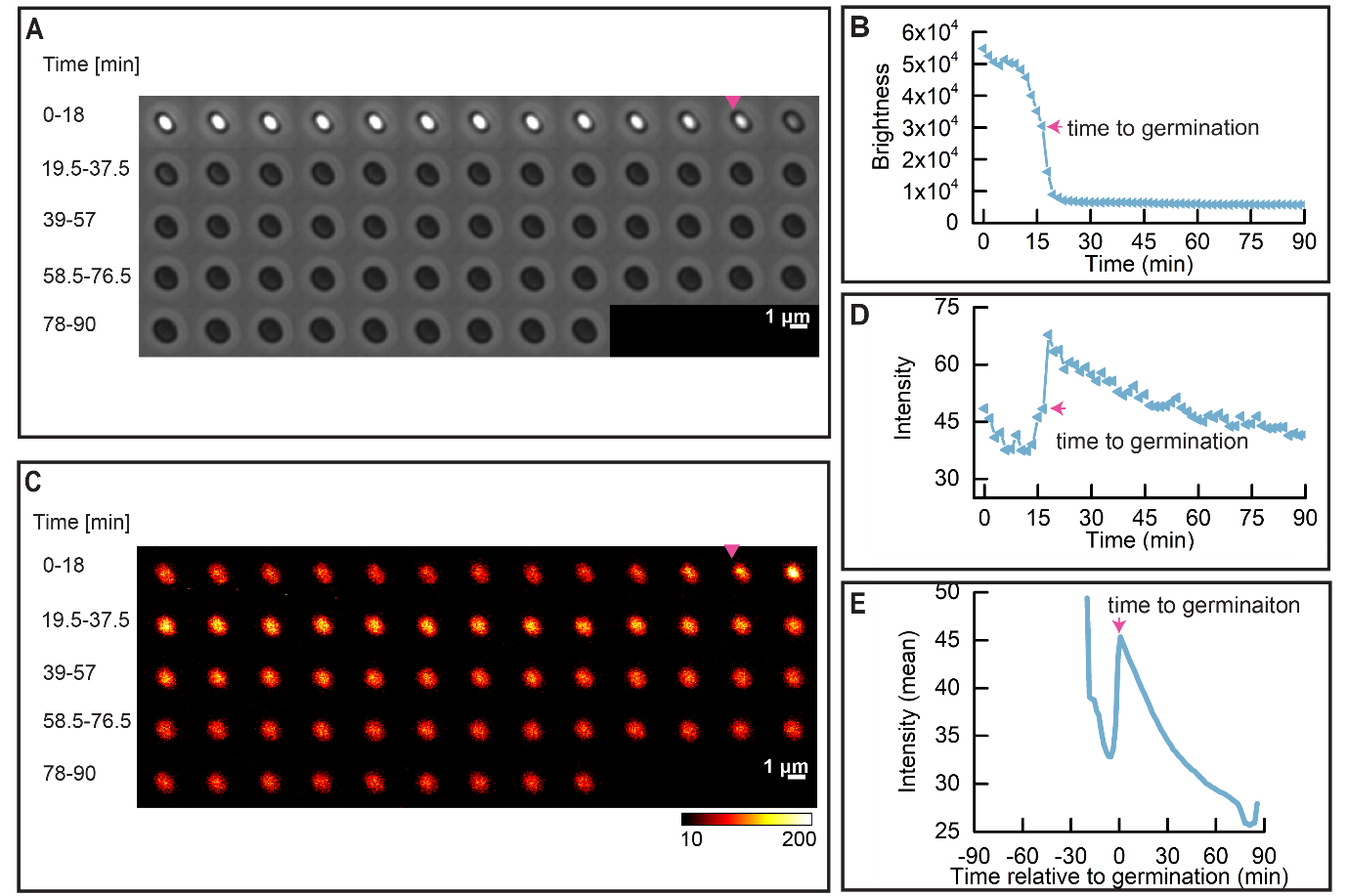

**Figure S2**. Dynamics of heat activated SpoVAEa-SGFP2 during spore germination. Spore germination was triggered by (10 mM each) AGFK after 5 hours of heat activation at 65 °C. (A) Phase contrast time lapse images of a single PS832 SpoVAEa-SGFP2 spore. (B) The brightness profile corresponding to the images shown in panel A. (C) The fluorescence time lapse images of the same spore shown in panel A. (D) The SpoVAEa-SGFP2 fluorescence intensity profile corresponding to images in the panel C. The magenta arrow indicates the ‘time to germination’. (E) Average of 562 synchronized single SpoVAEa-SGFP2 spore intensity traces. Synchronization defines t=0 min as the ‘time to germination’. 595 spores’ were tracked by microscope for 90min, 94.5% of them completed germination. Notably, the drop in SpoVAEa-SGFP2 fluorescence intensity before the ‘time to germination’ was most likely due to bleaching of the spore’s autofluorescence.

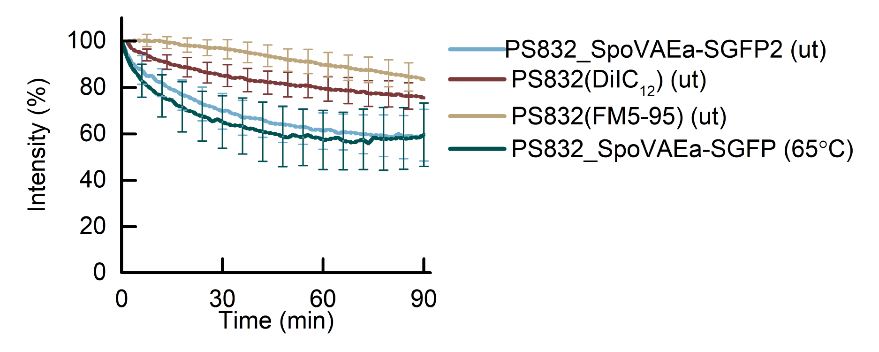

**Figure S3**. Loss of fluorescent intensity of dormant spores tracked by time-lapse imaging. Analysed dormant spores came from the same populations presented in Fig. 2, 3, 4, and S2, respectively, and these spores didn’t respond to AGFK induced germination in the 90 min time-lapse imaging process. The image acquisition and image analysis were detailed in the Materials and Methods, as well as legends of corresponding figures mentioned above. In total, the examined spore numbers of PS832_SpoVAEa-SGFP2 (ut), PS832(DiIC_12_) (ut), PS832(FM5-95) (ut), and PS832_SpoVAEa-SGFP2 (65°C) spores were 195, 119, 92, and 33, respectively. ut, untreated spores. 65°C, heat treatment at 65°C for 5 hours.

Supplementary Table 1. Pearson correlation between spore brightness and green autofluorescence or fluorescence intensity of SpoVAEa-SGFP2.

| T (°C) | **PS832** | | | **PS832 SpoVAEa-SGFP2** | | | **PS4150** | | | **PS4150 SpoVAEa-SGFP2** | | |
| --- | --- | --- | --- | --- | --- | --- | --- | --- | --- | --- | --- | --- |
|  | No. spores | Pearson correlation | | No. spores | Pearson correlation | | No. spores | Pearson correlation | | No. spores | Pearson correlation | |
|  |  | r | p-value |  | r | p-value |  | r | p-value |  | r | p-value |
| ut | 672 | 0.54 | < 0.01 | 1201 | -0.13 | < 0.01 | 700 | -0.021 | 0.57 | 1287 | 0.0068 | 0.81 |
| 40 | 1577 | 0.56 | < 0.01 | 1057 | 0.036 | 0.25 | 1135 | 0.083 | < 0.01 | 1254 | 0.083 | < 0.01 |
| 50 | 718 | 0.54 | < 0.01 | 948 | 0.1 | < 0.01 | 553 | -0.008 | 0.85 | 1174 | 0.041 | 0.17 |
| 60 | 1207 | 0.28 | < 0.01 | 951 | 0.19 | < 0.01 | 839 | 0.013 | 0.72 | 1511 | 0.033 | 0.20 |
| 65 | 1250 | 0.29 | < 0.01 | 1121 | 0.29 | < 0.01 | 1248 | 0.047 | 0.11 | 1249 | 0.017 | 0.56 |
| 70 | 1016 | 0.47 | < 0.01 | 1123 | 0.35 | < 0.01 | 1377 | 0.047 | 0.079 | 1358 | 0.023 | 0.39 |
| 75 | 816 | 0.32 | < 0.01 | 1133 | 0.32 | < 0.01 | 1541 | -0.18 | < 0.01 | 937 | 0.3 | < 0.01 |
| 80 | 444 | 0.37 | < 0.01 | 1273 | 0.045 | 0.11 | 1967 | 0.53 | < 0.01 | 1109 | 0.7 | < 0.01 |

ut: untreated spores

Supplementary Table 2. Supplementary Table 1. Pearson correlation between spore brightness and fluorescence intensity of dye stained IM.

| T (°C) | **832(DiIC_12_)** | | | **832(FM5-95)** | | |
| --- | --- | --- | --- | --- | --- | --- |
|  | No. spores | Pearson correlation | | No. spores | Pearson correlation | |
|  |  | r | p-value |  | r | p-value |
| ut | 876 | -0.012 | 0.73 | 1739 | 0.17 | < 0.01 |
| 40 | 955 | -0.012 | 0.53 | 1136 | 0.26 | < 0.01 |
| 50 | 1091 | -0.041 | 0.18 | 1383 | 0.2 | < 0.01 |
| 60 | 1335 | -0.044 | 0.11 | 1284 | 0.23 | < 0.01 |
| 65 | 612 | -0.075 | 0.064 | 1350 | 0.29 | < 0.01 |
| 70 | 583 | -0.013 | < 0.01 | 1639 | 0.54 | < 0.01 |
| 75 | 1472 | -0.22 | < 0.01 | 932 | 0.76 | < 0.01 |
| 80 | 1049 | -0.37 | < 0.01 | 2368 | 0.36 | < 0.01 |

ut: untreated spores
